# Leptin alleviates obesity hypoventilation *via* serotonergic pathways

**DOI:** 10.64898/2026.06.23.734005

**Authors:** Dashdulam Davaanyam, Melanie Alexis Ruiz, Junia L de Deus, Mi-Kyung Shin, Cole R Winston, Xin Wang, Mateus R. Amorim, David Mendelowitz, Vsevolod Y. Polotsky

**Affiliations:** Department of Anesthesiology and Critical Care Medicine, George Washington University School of Medicine and Health Science; Department of Pharmacology and Physiology, George Washington University School of Medicine and Health Science; Department of Physiology, Medical School of Ribeirão Preto, SP, Brazil

**Keywords:** leptin, serotonin, sleep apnea, hypoventilation, obesity

## Abstract

**Rationale:** There is no effective pharmacotherapy for obesity hypoventilation syndrome (OHS). Intranasal leptin augments the hypercapnic ventilatory response (HCVR), attenuates upper airway obstruction, and increases ventilation during sleep in diet-induced obese (DIO) mice. Respiratory effects of leptin can be attenuated by serotonergic antagonists.

**Objectives:** To establish if serotonergic innervation of the hypoglossal motoneurons (XII MN) mediates effects of leptin on OHS.

**Methods:** We examined effects of intranasal leptin on the HCVR, sleep architecture, arousal latency, flow limited (obstructed) and non-flow limited breathing, genioglossus muscle (GG) activity and metabolic rate across sleep/wake states in the presence and absence of serotonergic neurons innervating XII MN in DIO *Sert-flp* mice expressing *FlpO* recombinase in the serotonergic neurons. These mice were transfected into the XII MN with retrograde adeno-associated virus carrying either *FlpO*-dependent caspase or control yellow fluorescent protein (YFP).

**Measurements and Main Results:** Control YFP virus was densely localized to the serotonergic neurons of the medullary raphe (MR), but not the dorsal raphe (DR), and these neurons were ablated by caspase. Leptin enhanced the HCVR, increased arousal latency in males, but not in females, and these effects were abolished by caspase. Neither leptin nor caspase affected sleep architecture or metabolic rate. Leptin increased GG activity awake and during NREM sleep, attenuated pharyngeal obstruction and increased minute ventilation in NREM and REM sleep. All effects of leptin were abolished by the *FlpO*-dependent caspase.

**Conclusions:** Leptin treats OHS by stimulating MR serotonergic neurons, which project to XII MN and stimulate pharyngeal muscles during sleep.

## Introduction

Obstructive sleep apnea (OSA) is characterized by loss of upper airway muscle tone during sleep leading to recurrent periods of upper airway obstruction, intermittent hypoxia and hypercapnia^1^. Obesity hypoventilation syndrome (OHS) is a life-threatening severe form of sleep disordered breathing (SDB) characterized by daytime hypercapnia and hypoventilation during sleep. OHS is observed in 10-20% of obese patients with OSA^2,3^. OHS is driven by decreased pH/CO_2_ sensitivity and depressed ventilatory responses to CO_2_^2,4–6^, which is a consequence of recurrent hypercapnia in severe OSA^7^.

Neuromuscular mechanisms play an important role in maintaining upper airway patency during sleep^8–11^, especially the hypoglossal motoneurons (XII MN)^12–15^ and hypoglossal nerve^16–18^ innervating tongue muscles including the main pharyngeal dilator, genioglossus muscle (GG)^11,17,19–23^. Pharmacological approaches focused on adrenergic, cholinergic and serotonergic mechanisms have been tested in animal models and patients with OSA^24–29^, but not in patients with OHS.

Our efforts are focused on leptin, an adipose-produced hormone, which suppresses food intake and increases metabolic rate^30–33^. In leptin deficient obese *ob/ob* mice, leptin augments hypercapnic sensitivity^34,35^ and improves upper airway patency during sleep^36–38^. However, human obesity is characterized by high levels of circulating leptin and resistance to metabolic^39,40^ and respiratory effects^41,42^ of systemic leptin. We have previously shown that diet-induced obese (DIO) mice emulate all features of OHS, including awake hypercapnia, suppressed hypercapnic ventilatory response (HCVR), inspiratory flow limitation with hypoventilation during sleep, high plasma leptin levels and leptin resistance^43–46^. Resistance to intravenous and intraperitoneal leptin treatment in DIO mice can be overcome by intranasal (IN) leptin administration^43,47^, which increased minute ventilation during sleep and attenuated inspiratory flow limitation, a hallmark mark manifestation of pharyngeal obstruction in OSA^43^. We identified neurons expressing a long active isoform of leptin receptor (LEPR^b^+) in the dorsomedial hypothalamus (DMH) as a likely target for the respiratory responses to leptin^48,49^. Furthermore, we showed that 5-HT receptor blockers and elimination of LEPR^b^+ DMH neurons projecting to the 5-HT dorsal raphe (DR) nucleus abolished effects of leptin on HCVR and upper airway patency during sleep^49^. Nevertheless, the mechanisms by which leptin treats SDB are not fully understood. Leptin decreased pharyngeal collapsibility under anesthesia^38^, but effects of leptin on GG muscle tone during sleep have not been studied. The role of serotonergic neurons in leptin-induced HCVR augmentation and pharyngeal patency during sleep has not been elucidated. Finally, the effect of the leptin-5-HT axis on sleep architecture has not been studied.

We hypothesized that leptin’s therapeutic effects in SDB are modulated by 5-HT neurons projecting to hypoglossal motoneurons, which innervate GG, and that the elimination of these neurons abolish leptin effects on GG activity, HCVR and SDB. We examined our hypothesis in *Sert-flp* mice, which express codon-optimized *FlpO* recombinase under control of the mouse *Slc6a4* (solute carrier family 6 serotonin transporter, member 4) and enable to excise FRT-flanked sequences in 5-HT-expressing cells^50^. We transfected DIO *Sert-flp* mice with retrograde AAV expressing *FlpO*-dependent caspase or control virus into the hypoglossal nucleus and examined effects of IN leptin on the HCVR, arousal latency, sleep architecture, GG activity and breathing during sleep in the presence or absence 5-HT neurons projecting to the XII MN.

## Methods

### Animals

All experimental protocols were approved by the Institutional Animal Care and Use Committee (IACUC) at The George Washington University (Protocol No. A2025-152, A2025-155 and A2023-009). The study has been conducted in accordance to the American Physiological Society’s (APS) guidelines for the care and use of laboratory animals^51^ and the Animal Research Reporting In Vivo Experiments (ARRIVE) guidelines^52^.

In total, 44 age matched (26-30 weeks) *Sert-Flp* mice (B6. Cg -*Slc6a4^tm1.1(flop)Luo^*/J, JAX #034050) from the Jackson Laboratory (Bar Harbor, ME, USA) were used in this study, including 31 males and 13 females. Male *Lepr^b^ -Cre-* mice (B6.129 (Cg) LEPR^tm (cre)Rck^/J; JAX #008320, n = 3) were also from the Jackson Laboratory. Water and food were available ad *libitum*. Mice were fed with a high fat diet containing 60kcal% fat (D12492, Research Diets Inc., New Brunswick, NJ, USA). Mice were housed at thermoneutral conditions (28-30°C) and the 12-hours light/dark cycle. All animals were acclimated to the housing conditions prior to the initiation of experiments for at least 1 week. At the completion of experiments, mice were euthanized by anesthetic overdose and cervical dislocation.

### Viral vector administration

Viral vector administration were carried out as described previously^53^ with minor modifications. All surgical procedures were performed under sterile, aseptic conditions. Mice were anesthetized with isoflurane (2-3% for induction in closed chamber; maintained at 1-2% during surgery) and positioned in a stereotaxic apparatus (Kopf Instruments, Tujunga, CA, USA). The depth of anesthesia was confirmed by absence of withdrawal reflex following tail pinch. Body temperature was maintained throughout the procedure using a heating pad. *FlpO*-dependent retrograde control viral vector pAAV-Ef1a-fDIO-EYFP (#55641; ≥7x10^12^ vg.mL^-1^; Addgene, Watertown, MA, USA) virus or retrograde *FlpO*-dependent caspase virus, AAV2-retro.EF1a.FlexFrt.taCasp3.T2A.TEVp.WPRE.hGH (≥7x10^12^ vg.mL^-1^; Research Vector Core at Franklin Biolabs, King of Prussia, PA, USA) were delivered using disposable glass micropipettes (VWR Int., Radnor, PA, USA) into the hypoglossal nucleus using the following stereotactic coordinates from the bregma: 7.20 mm anterior-posterior, 0.20 mm medial-lateral and 5.0 mm dorsal-ventral. AAV1-EF1a-double floxed-hChR2 (H134R)-EYFP-WPRE-HGHpA (#20298-AAV1, ≥7x10^12^ vg.mL^-1^; Addgene) was injected in the dorsomedial hypothalamus (DMH) of *Lepr^b^-Cre* mice. The stereotactic coordinates were determined based on the mouse brain atlas^54^. Viral infusion was performed slowly to minimize tissue damage and allow diffusion within the target area. Following surgery, mice received buprenorphine XR (Ethiqa; 3.25 mg/kg^-1^, subcutaneous) for postoperative analgesia and were monitored daily during recovery.

### Intranasal leptin administration

Mice were randomized to receive leptin or vehicle treatments in a crossover experimental design, with successive administrations separated by 5-7 days. Intranasal leptin delivery was performed as previously described^55,56^ with minor modifications. Mice were anesthetized with isoflurane (1-2% for induction and maintenance) and positioned supine at a 90°C angle for intranasal administration. Recombinant leptin (#498-OB; 0.8 mg/kg^-1^; R&D Systems, Minneapolis, MN, USA) was prepared in a total volume of 24 µL consisting of 1% bovine serum albumin (BSA) in phosphate-buffered saline (PBS). Vehicle-treated mice received 1% BSA containing PBS. Leptin or vehicle solution were administered intranasally using pipette tips (TipOne; USA Scientific, Ocala, FL, USA) in alternating 3 µL drops delivered into each nostril, for a total 12 µL per nostril administered in two sequential doses. Following each application, mice were maintained in the supine position for approximately 30-40 seconds to facilitate nasal absorption and minimize drainage. Animals were subsequently returned to the prone position and monitored until recovery from anesthesia.

### Surgical Implantation of EEG and EMG Electrodes

Head mount implantation with electroencephalogram (EEG) and electromyogram (EMG) electrodes for sleep recordings was performed as previously described^57–62^. All surgical procedures were conducted under sterile and aseptic conditions. Mice were weighed and anesthetized with 1-2% isoflurane and positioned in the stereotaxic system (Kopf Instruments, Tunjunga, CA, USA). After making a longitudinal midline incision to exposure the skull, the surgical site was disinfected with providone-iodine (Betadine) solution, and the underlying fascia was gently cleared from the skull surface. A four-pin EEG/EMG head mount (No.8201-SS, Pinnacle Technology, Lawrence, KS, USA) was placed over top of bregma. Two silver electrodes of 0.1mm (No.8209, Pinnacle Technology) and two of 0.12mm (No.8212, Pinnacle Technology) were screwed through the holes and covered with silver conductive epoxy (No.8331, MG Chemicals, ON, Canada) to provide unipolar conductive EEG electrodes. For EMG electrodes, insulted leads were tunneled subcutaneously and placed bilaterally over the nuchal muscle posteriorly to the skull. The head mount assembly was fixed to the skull using dental acrylic (Lang Dental, IL, USA). Upon surgical completion, mice were provided with subcutaneous buprenorphine XR (3.25 mg/kg^-1^) and placed a thermal heat source during anesthetic recovery. Animals were allowed to recover for a minimum of 1 week before the initiation of sleep recording experiments.

### Whole-Body Plethysmography and Respiratory Analysis

Mice were placed in a modified whole-body plethysmography chamber system (WBP; EMMS, Hampshire, UK) for respiratory measurements, as previously described^57–62^. A two-stage crossover study design was used. Mice were acclimated to the WBP chambers for least 1 day before the sleep studies. Recordings were performed under thermoneutral condition of 28-30°C and 90% humidity for 6 hours (from 10:00 AM to 4:00 PM). Mouse weight and rectal temperature were measured at the beginning and end of the study. All signals were digitized at 1000 Hz and recorded in LabChart 7 Pro (ADInstruments, Sydney, Australia). The Drorbaugh and Fenn equation was used to calculate the tidal volume signal from chamber pressure^63^. Respiratory analysis was performed as previously described ^57,61^. Minute ventilation (V_E_) was assessed across the entire sleep recording. Custom software was used to demarcate the start and end of inspiration and expiration for subsequent calculations of timing and amplitude parameters for each respiratory cycle. The instantaneous respiratory rate (RR, breaths/min) was calculated as the reciprocal of the respiratory period, and the instantaneous V_E_ (mL/min) was product of the RR and VT for each breath. We then utilized the airflow and respiratory effort signals to develop an algorithm for detecting upper airway obstruction during sleep. Obstruction was characterized by the development of inspiratory flow limitation (IFL), which was defined by an early peak of inspiratory flow followed by a plateau or by the late inspiratory peak in airflow exceeding the early peak^57^. Apneas were defined as ≥90% drops in the flow channel during sleep for at least two breath cycles or ≥0.7 s regardless of the presence of arousals. Sighs were defined by breaths with > 2-fold increase in the tidal volume compared to average. The apnea and sigh indexes were calculated by dividing the number of apneas by the total sleep time (in hours)^58^.

### Sleep scoring

Studies were manually scored for sleep-wake stages in 5-second epochs using established EEG and EMG criteria as previously described^64^. Wakefulness was identified by high-frequency (10-20 Hz) and low-amplitude EEG signals. NREM sleep featured high-amplitude and low-frequency (2-5 Hz) in the EEG. REM sleep was scored as low-amplitude mixed frequencies (5-10 Hz) in the EEG and EMG muscle atonia. Sleep efficiency was calculated as the total sleep time divided by the total recording time after sleep onset. Arousals from sleep were manually scored as abrupt increases in the frequency of EEG signals for at least 0.5 seconds and less than 2.5 seconds (<50% of each sleep epoch). Arousals were marked in one of the EEG channels when they were preceded by at least 2.5 seconds of stable sleep. In REM sleep, EEG arousals were scored when associated with increases in EMG activity. Arousal index was calculated as the total number of arousals divided by the total sleep time in hours^64^.

### Arousal scoring

Arousals from sleep were manually scored as previously described^64^. Arousals were defined as abrupt increases in EEG frequency lasting between 0.5 and 2.5 seconds. Each event was required to be preceded by at least 2.5 seconds of stable breathing. Arousal index was calculated as the total number of arousals divided by the total sleep time.

### Hypercapnic ventilatory response (HCVR)

Mice were acclimated to the WBP (EMMS) chamber prior to experiments, in accordance with procedures previously established by our group^65,66^. All measurements were performed under thermoneutral condition (28-30°C) during the light phase. Mice were acclimated with a continuous-bias flow controlled with mass flow controllers in room air for 30 minutes. They were exposed to a gas mixture of 8% CO_2_, 21% O_2_, and balanced in nitrogen. For exposure, room air was switch to the hypercapnic mixture, and analysis were done after 1 min of exposure when the ventilation reached a plateau. Tidal volume (V_T_), respiratory rate (RR), and minute ventilation (V_E_) were measured in mice at baseline (room air) and hypercapnic ventilatory response (HCVR) was determined in each animal by the slope of the relationship between minute ventilation (V_E_) and inspired CO2 (0-8%) during wakefulness via linear least-squares regression analysis.

### Respiratory arousal threshold (arousal latency)

Respiratory arousal threshold was assessed as previously described^67,68^. All recordings were performed in a WBP chamber system while mice were simultaneously monitored for sleep-wake state. Gas concentrations inside the WBP chamber were continuously controlled using an automated CO_2_/O_2_ gas mixer (CWE, Ardmore, PA, USA). Experiments were conducted during the light phase between 1:00 PM and 4:00 PM. Mice were exposed to repeated cycles of normoxia and hypercapnia consisting of 5 minutes of normoxia followed by 40 seconds of 8% CO_2_, 21% O_2_, balanced in N_2_. Respiratory arousal threshold was determined by measuring the latency from the onset of 8% CO_2_ exposure to EEG-defined arousal from NREM sleep. Only trials in which mice maintained stable NREM sleep for at least 30 seconds immediately before CO_2_ exposure were included in the analysis. Arousal latency was calculated as the time required for mice to transition from NREM sleep to wakefulness within the 40 seconds CO_2_ exposure period.

### Metabolic measurements

Metabolic studies were performed as previously described^56,60,66^.Mice were placed in the Promethion Core Mouse Metabolic System (SABLE Systems Int., North Las Vegas, NV, USA) cages for 48-72 hours acclimation period followed by 24 hours of continuous recordings starting at 10:00 AM. The experiments were done in a randomized crossover design throughout the light phase and dark phase. The SABLE cages are sealed and equipped with O_2_ electrochemical sensors, CO_2_ infrared sensors, and infrared beam movement sensors. Every 5 minutes, consumed O_2_ (VO_2_), and produced CO_2_ (VCO_2_) were collected, and measurement were utilized to calculate the respiratory exchange ratio (RER). An array of infrared photo beams that surrounded the metabolic cage was quantified by the counts of infrared beam interruptions. Total horizontal and vertical beam breaks were summed and presented as motor activity. Mice had free access to food and water during the measurements. Metabolic cage kept on a 12-hours light/12-hours dark cycle (7:00 AM-7:00 PM lights on/lights off) with food and water ad *libitum* and a consistent environmental temperature of 29-30°C.

### Electromyography of the Genioglossus muscle (EMG_GG_)

Electromyographic activity of the genioglossus (GG) muscle was recorded as previously described^25,69^ with minor modifications. Mice were anesthetized with isoflurane (2-3% for induction; maintained at 1-2% during surgery). A custom head mount containing extended GG electrodes was implanted using procedures similar to those used for EEG/EMG sleep recordings. The extended GG electrodes were tunneled subcutaneously from the scalp incision to a mandibular incision beneath the tongue. Mice were positioned supine under analgesia, and two Teflon-insulated stainless steel hook electrodes (A-M Systems, Carlsborg, WA, USA) were inserted into the genioglossus muscle between the base of the mandible and the exit point of the lingual vein and nerve bundle. Electrodes were inserted parallel to the muscle fibers to a depth of approximately 3 mm and secured with surgical thread. Excess wire exposed outside the muscle was trimmed and removed before closure of the incision. Following surgery, mice recovered for 5-7 days before initiation of recording experiments. GG activity and sleep-wake states were continuously recorded and manually scored and established EEG and EMG criteria as previously described. Upon completion of recordings, mice were monitored during recovery and received buprenorphine (3.25 mg/kg^-1^, subcutaneous) for postoperative analgesia.

### Immunofluorescence

For the immunofluorescent analysis, mice were anesthetized with isoflurane (1-2%) and transcardially perfused with ice-cold phosphate-buffered saline (PBS; 0.01M, pH 7.4), followed immediately by 4% paraformaldehyde (PFA) prepared in PBS. Brains were carefully removed and postfixed in 4% PFA at 4°C for 48 hours. Brains were sectioned coronally at 30 µm thickness using a vibratome (Leica VT1000S; Leica Microsystems, Wetzlar, Germany), and free-floating sections were collected in PBS for immunohistochemical processing. Sections were the incubated in blocking solution containing 10% normal goat serum (NGS), 1% bovine serum albumin (BSA), and 0.1% Triton X-100 in 0.01M PBS for 1 hour at room temperature to reduce nonspecific binding and permeabilize tissue. Free-floating sections were then incubated with primary antibody against the serotonergic neuronal marker tryptophan hydroxylase-2 (rabbit anti-TPH2; 1:500; Novus Biologicals, Centennial, CO, USA and rat anti-TPH, 1:200; Abcam, Cambridge, MA, USA) overnight at 4 °C. Following primary antibody incubation, sections were washed thoroughly in PBS and incubated for 1 hour at room temperature with Alexa Fluor 647-conjugated goat anti-rabbit IgG secondary antibody (1:500; Invitrogen, Carlsbad, CA, USA) and Alexa Fluor 647-conjugated goat anti-rat IgG secondary antibody (1:500; Invitrogen) together with DAPI nuclear stain (4,6-diamidino-2-phenylindole; 1:500; Invitrogen). Sections were then mounted onto glass slides and cover slipped using Prolong Gold antifade mounting medium (Invitrogen). Fluorescent images were acquired using a Zeiss LSM 980 confocal microscope (Carl Zeiss Microscopy GmbH, Jena, Germany).

### Statistical Analysis

Results were processed and analyzed using Prism.11 (GraphPad Software Inc., San Diego, CA, USA). Wilcoxon matched-pairs signed was used to compare differences within subjects. Mann Whitney test was used to compare differences between the groups. Correlations between variables were analyzed using two-tailed Spearman’s rank correlation test. Data were analyzed using a two-way mixed-effects model with leptin treatment and caspase ablation as fixed factors. Interaction effects between leptin and caspase were assessed. When significant effect was detected, post hoc multiple- comparison tests were performed. Graphically, data were plotted using boxplots [median ±1.5*interquartile range (IQR)] showing individual mice. Statistical significance was defined as p<0.05.

## Results

### Visualization of 5-HT neurons projecting to the hypoglossal nucleus using retrograde AAV in *Sert-Flp* mice and the effect of *FlpO*-dependent caspase

We deleted 5-HT-expressing neurons projecting to the hypoglossal motor neurons (XII MN) using retrograde AAV-caspase and compared the results with retrograde AAV-YFP labeling controls. Schematic representations of AAV-retro-YFP (**Figure 1A**) and AAV-retro caspase (**Figure 1B**) administration into the XII MN illustrated viral spread (YFP-green) to the 5-HT neurons (TPH-red), with colocalization shown in orange.

**Figure 1.**
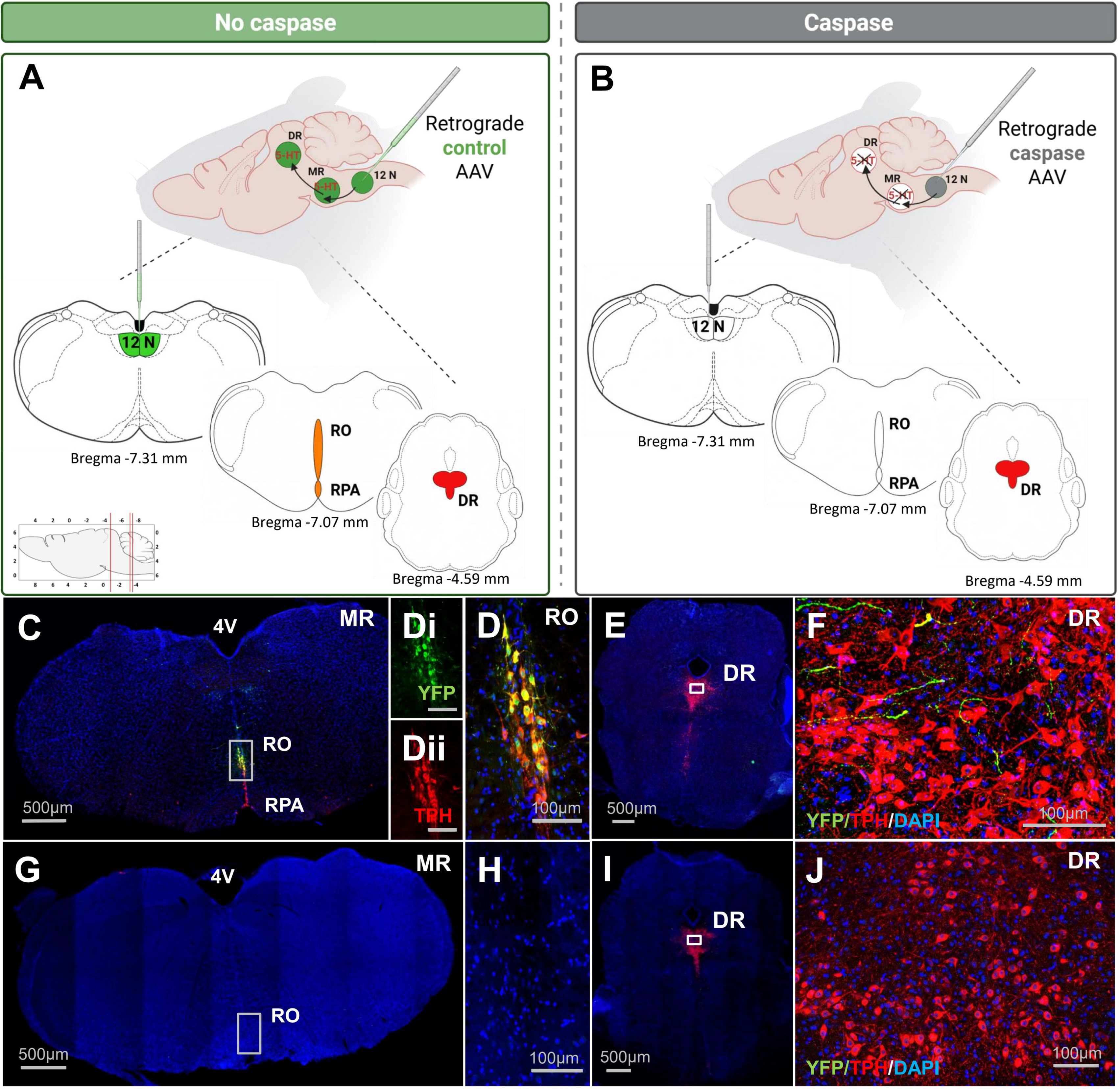
Transfection of the hypoglossal nucleus (12 N) with the retrograde AAV carrying *Flp*O-dependent YFP (A, C-F) or caspase (B, G-J) in *Sert-flp* mice. Schematic presentation of **(A)** AAV retro-YFP and **(B)** AAV retro-caspase administration and the viral spread (green) in relationship to the serotonergic (5-HT+) neurons (red); colocalization is shown in orange, and the elimination of 5-HT+ neurons is illustrated by the disappearance of red color. **(C)** Retrograde YFP labeling (green) was present in 5-HT+ (tryptophan hydroxylase, TPH; red) neurons of the raphe obscurus (RO) following control retrograde AAV transfection of the XII MN. The outlined region is enlarged in **(D-Dii)** to show **(D)** retrograde YFP labeling of the RO neurons after XII MN administration. **(E)** TPH-positive serotonergic neurons (red) in the DR were surrounded by green fibers arising from the 5-HT neurons directly projecting to the XII MN following control retrograde AAV-YFP transfections into the XII MN; the outlined area is enlarged in **(F)**. **(G)** Retrograde AAV caspase administered to XII MN eliminated all serotonergic neurons in the RO; the outlined region is enlarged in **(H)**. **(I)** TPH-positive serotonergic neurons (red) remained in the DR after caspase transfection of the XII MN; the outlined region is enlarged in **(J)**. 4V, 4^th^ ventricle. Scale bars 500 µm; 100 µm (enlarged view)

Retrograde AAV-YFP control virus transfection into the XII MN produced robust YFP labeling in tryptophan hydroxylase (TPH) positive 5-HT neurons within the raphe obscurus (RO) and raphe pallidus (RPA) (**Figure S1**) subdivisions of the medullary raphe (MR), showing direct serotonergic projections from the MR to the XII MN (**Figure 1C, D**). In the DR, TPH positive serotonergic neurons were surrounded by YFP positive fibers following control retrograde AAV-YFP administration in the XII MN (**Figure 1E, F**), suggesting serotonergic projections from the neurons directly innervating XII MN. Retrograde AAV-caspase administration into XII MN eliminated serotonergic neurons in the RO **(Figure 1G, H**) and RPA, which was evident from the absence of TPH positive neurons in MR. In contrast, TPH positive neurons in the DR remained following caspase transfection of the XII MN (**Figure 1I and J**), indicating the lack of direct projections from DR to XII MN.

### The effect of intranasal leptin on the HCVR and hypercapnic arousal latency and the role of the 5-HT neurons projecting to the XII MN

We tested whether intranasal leptin affects ventilatory and arousal responses to CO_2_ through 5-HT neurons projecting to the hypoglossal nucleus in male and female *Sert-Flp* mice. There was no effect of leptin or caspase on body weight (48.0 ± 0.6 g in males and 49.8 g ± 1.8 g in females of all experimental groups) or body temperature (36.3 ± 0.05 °C in males and 36.2 ± 0.12°C in females of all experimental groups).

Hypercapnia rapidly elicited arousals in sleeping animals, therefore the HCVR was tested only during wakefulness. Under room air conditions, minute ventilation (V_E_), tidal volume (V_T_), and respiratory rate (RR) were not significantly different between groups (**Figure 2A-C**). During 8% CO_2_ exposure, leptin significantly elevated V_E_ (p=0.02; **Figure 2D**) and V_T_ (p=0.04; **Figure 2E**) in control transfected male mice, while RR (**Figure 2F**) was unaffected. Consequently, leptin significantly enhanced the HCVR (p=0.004; **Figure 2G**) in control virus transfected male mice. Caspase-induced ablation of XII-projecting 5-HT neurons abolished the leptin on HCVR. In female *Sert-Flp* mice, intranasal leptin did not significantly alter baseline or ventilatory response to 8% CO_2_ exposure, regardless of the virus treatment (**Figure 2H-O**).

**Figure 2.**
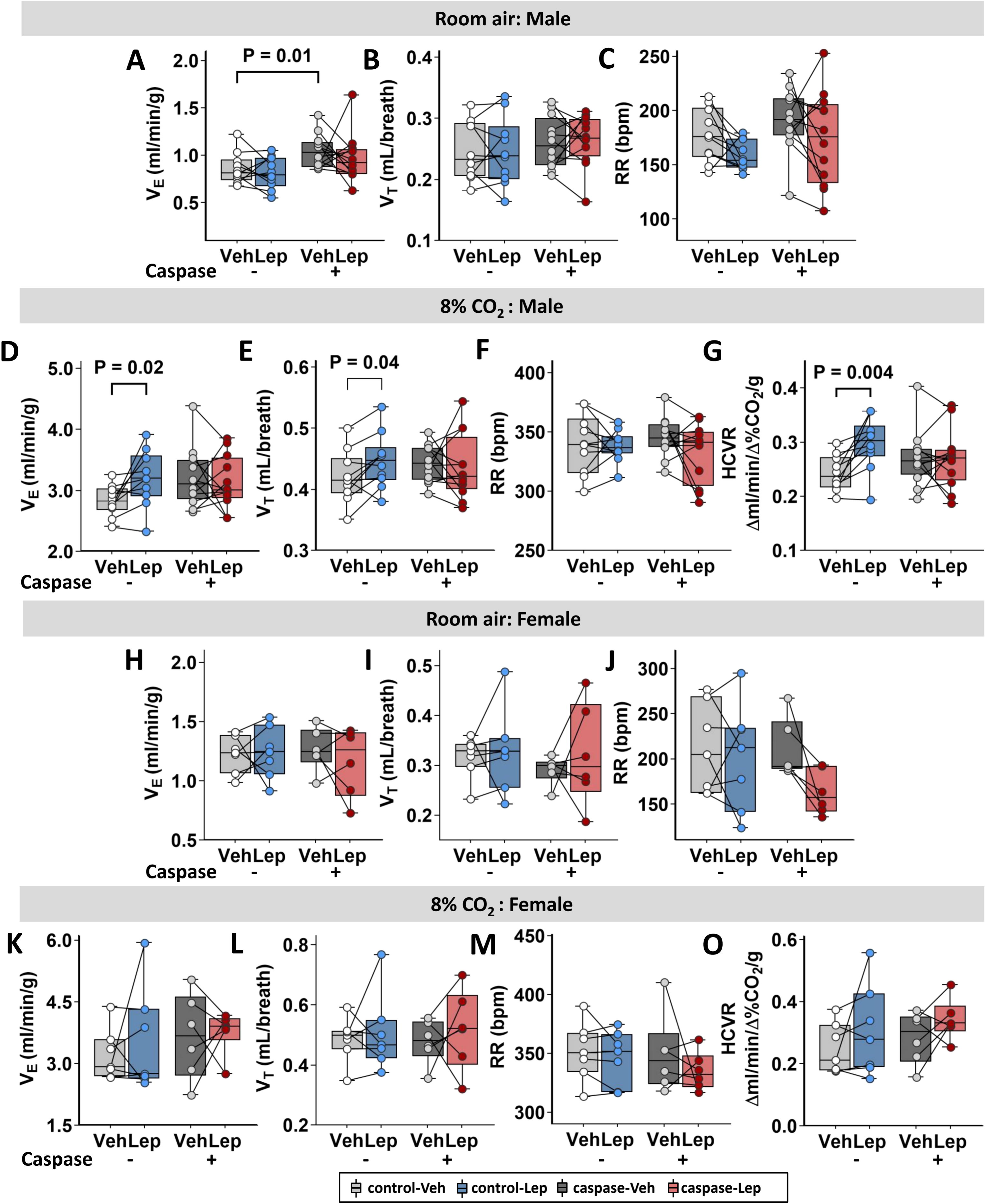
The effect of intranasal leptin on the hypercapnic ventilatory response (HCVR). Intranasal leptin (Lep) augmented minute ventilation (V_E_) during 8% CO_2_ exposure compared to vehicle (Veh) and enhanced the HCVR in male, but not in female *Sert-Flp* mice transfected with the control virus. Lep-induced enhancement of ventilation was abolished by elimination of 5-HT+ neurons projecting to the hypoglossal nucleus. **(A-C)** Male mice under room air conditions: **(A)** Minute ventilation (V_E_), **(B)** tidal volume (V_T_) and **(C)** respiratory rate (RR). **(D-G)** Male mice during 8% CO_2_ exposure: **(D)** V_E_, **(E)** V_T_, **(F)** RR, and **(G)** HCVR. **(H-J)** Female mice under room air condition: **(H)** V_E_, **(I)** V_T_, and **(J)** RR. **(K-O)** Female mice during 8% CO_2_ exposure: **(K)** V_E_, **(L)** V_T_, **(M)** RR, and **(O)** HCVR. Male control virus, n=10; male caspase virus, n=12; female control virus, n=7; female caspase virus, n=6. Data are plotted using boxplots (median ± 1.5*interquartile range). Statistical analyses were performed using the Wilcoxon matched-pairs signed rank test or Mann-Whitney test. Exact P values are shown in the figures.

A representative recording of the hypercapnic arousal from NREM sleep is shown in **Figure 3A**. In male mice, transfected with the control virus, intranasal leptin trended to increase arousal latency in response to the CO_2_ challenge (p=0.06; **Figure 3B**). Furthermore, after leptin challenge male mice stayed asleep in the hypercapnic environment longer and require a higher CO_2_ level to wake up (**Figure 3C**). Elimination of 5-HT MR neurons projecting to the XII MN abolished this effect. These results indicate that intranasal leptin enhanced the HCVR and increased CO_2_ arousal latency in male mice through 5-HT neurons projecting to the XII MN, whereas female mice did not respond to leptin (**Figure 3D and E).**

**Figure 3.**
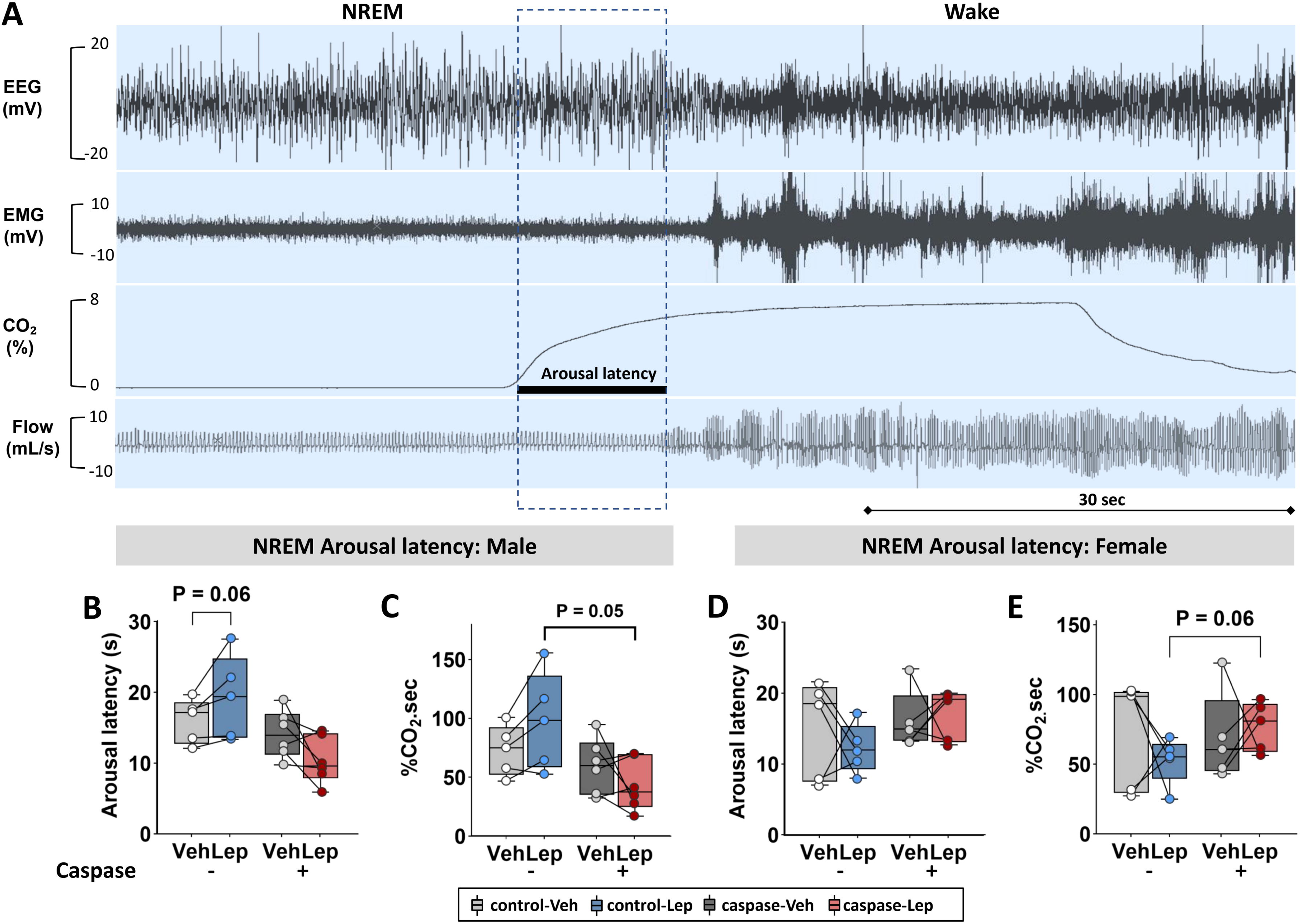
The effect of intranasal leptin on hypercapnic arousal latency. The deletion of 5-HT neurons projecting to XII MN altered NREM sleep arousal latency in a sex-dependent manner. **(A)** Representative screenshot of sleep recording during hypercapnic arousal testing. **(B)** Arousal latency in male mice. **(C)** Arousal latency x inspired CO_2_ concentration in male mice. **(D)** Arousal latency in female mice. **(E)** Arousal latency x inspired CO_2_ concentration in female mice Male control virus, n=5; male caspase virus, n=6; female control virus, n=5; female caspase virus, n=5. Data are plotted using boxplots (median ± 1.5*interquartile range). Statistical analyses were performed using the Wilcoxon matched-pairs signed rank test or Mann-Whitney test. Exact P values are shown in the figures. NREM: non-rapid eye movement sleep.

### The effect of intranasal leptin on metabolic parameters and sleep architecture and the role of the 5-HT neurons projecting to the XII MN

Given the lack of an effect of leptin on respiratory control in our current and previous experiments^49,70^, we proceeded with sleep studies only in male mice.

Neither intranasal leptin nor caspase-induced elimination of 5-HT neurons projecting to XII MN had any effect on body weight, body temperature, oxygen consumption, CO_2_ production, respiratory quotient, motor activity, sleep efficiency, sleep latency, the total amount of NREM and REM sleep, the number and duration of NREM and REM sleep bouts (**Table 1, Figure S4**). Leptin did not affect delta power of NREM, whereas caspase appeared to decrease it (**Figure S2**). In control mice, leptin significantly decreased the spontaneous arousal index (p=0.03; **Figure 4)**. After removal of the 5-HT neurons projecting to the XII MN leptin had an opposite effect increasing the arousal index (**Figure 4**, p < 0.01 for leptin x caspase interaction). Thus, intranasal leptin did not affect sleep macroarchitecture, but it decreased the number of arousals from sleep acting *via* 5-HT neurons innervating the XII MN.

**Figure 4.**
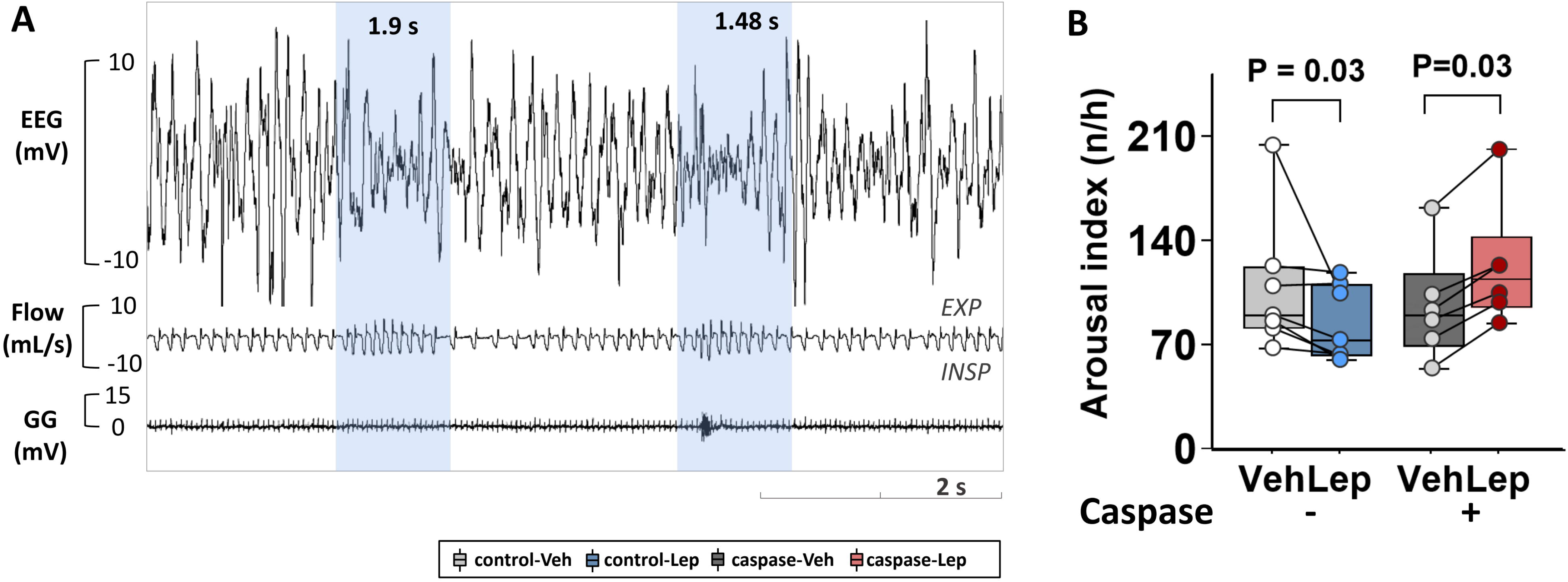
The effect of intranasal leptin on spontaneous arousals from sleep. **(A)** Representative screenshot of two spontaneous arousals from NREM sleep (shaded); **(B)** Arousal index (events per hour). Veh, vehicle; Lep, leptin. p < 0.01 for leptin x caspase interaction. Data are plotted using boxplots (median ± 1.5*interquartile range). Statistical analyses were performed using the Wilcoxon matched-pairs signed rank test or Mann-Whitney test. Two-way ANOVA revealed a significant interaction between leptin x caspase ablation. Exact P values are shown in the figures.

**Table 1.** Sleep architecture of diet-induced obese *Sert-flp* mice transfected with retrograde Flp dependent control or caspase AAV to the XII N.

| Groups | Sleep Efficiency (%) | Sleep (min) |  |  | Bouts |  |  |  | (%) |  | WASO (min) | Sleep Latency (min) |
| --- | --- | --- | --- | --- | --- | --- | --- | --- | --- | --- | --- | --- |
|  |  |  |  |  | Number |  | Average Length (min) |  |  |  |  |  |
|  |  | Total | NREM | REM | NREM | REM | NREM | REM | NREM | REM |  |  |
| Control-Veh | 32 ± 12.9 | 110.9 ± 47.8 | 94.4 ± 38.9 | 15.3 ± 12.3 | 152.9 ± 51 | 17.2 ± 13.4 | 0.6 ± 0.2 | 0.9 ± 0.2 | 85.6 ± 7.4 | 13.9 ± 7.5 | 193 ± 59.5 | 70.2 ± 41.8 |
| Control-Lep | 35.8 ± 11.5 | 123.6 ± 42.3 | 105.3 ± 34.2 | 19.7 ± 14.2 | 156.3 ± 37.5 | 20.9 ± 12.3 | 0.7 ± 0.2 | 0.9 ± 0.3 | 86.2 ± 8.4 | 14.8 ± 7.7 | 184.6 ± 29.2 | 61.4 ± 42.1 |
| Caspase-Veh | 35.2 ± 12.7 | 126.1 ± 51.9 | 103.5 ± 43.6 | 22.8 ± 13.1 | 187.6 ± 65.9 | 27.1 ± 15 | 0.6 ± 0.3 | 0.8 ± 0.2 | 82.5 ± 8.9 | 17.2 ± 8.9 | 206.9 ± 50.5 | 32.2 ± 29.2 |
| Caspase-Lep | 31.3 ± 9.7 | 105.3 ± 44.5 | 91.7 ± 39 | 16.2 ± 9 | 178.7 ± 66.3 | 20.8 ± 13.6 | 0.5 ± 0.1 | 1 ± 0.4 | 87.6 ± 7.9 | 14.7 ± 6.2 | 191.4 ± 49.4 | 44.9 ± 40.9 |

\*Values are presented as mean ± SD. Differences between groups were analyzed using a two-sample Student's t-test.

### The effect of intranasal leptin on breathing during NREM sleep and the role of the 5-HT neurons projecting to the XII MN

In control mice, intranasal leptin increased maximum inspiratory flow (V_I max_), minute ventilation (V_E_), and tidal volume (V_T_) during NREM sleep (**Figure 5A-D**). The respiratory analysis has been performed separately in inspiratory flow limited breaths identifiable by an early plateau in inspiratory flow (**Fig. 5F**), a characteristic of upper airway obstruction, in non-flow limited breaths (**Fig. 5K**). The percentage of inspiratory flow limited breath was < 5% and not significantly affected by leptin (**Fig. 5E**). Intranasal leptin was equally potent as a respiratory stimulant during both flow limited and non-limited breathing (**Figs. 5G-J, L-O**). Deletion of 5-HT neurons projecting to the XII MN prevented the effects of leptin on V_I max_, V_E_ and V_T_ during both flow limited and non-flow limited breathing, and revealed the leptin-induced decrease in the frequency of flow limited breaths. Taken together, caspase experiments show that 5-HT neurons projecting to the XII MN mediate effects of leptin on ventilatory control and upper airway patency during NREM sleep.

**Figure 5.**
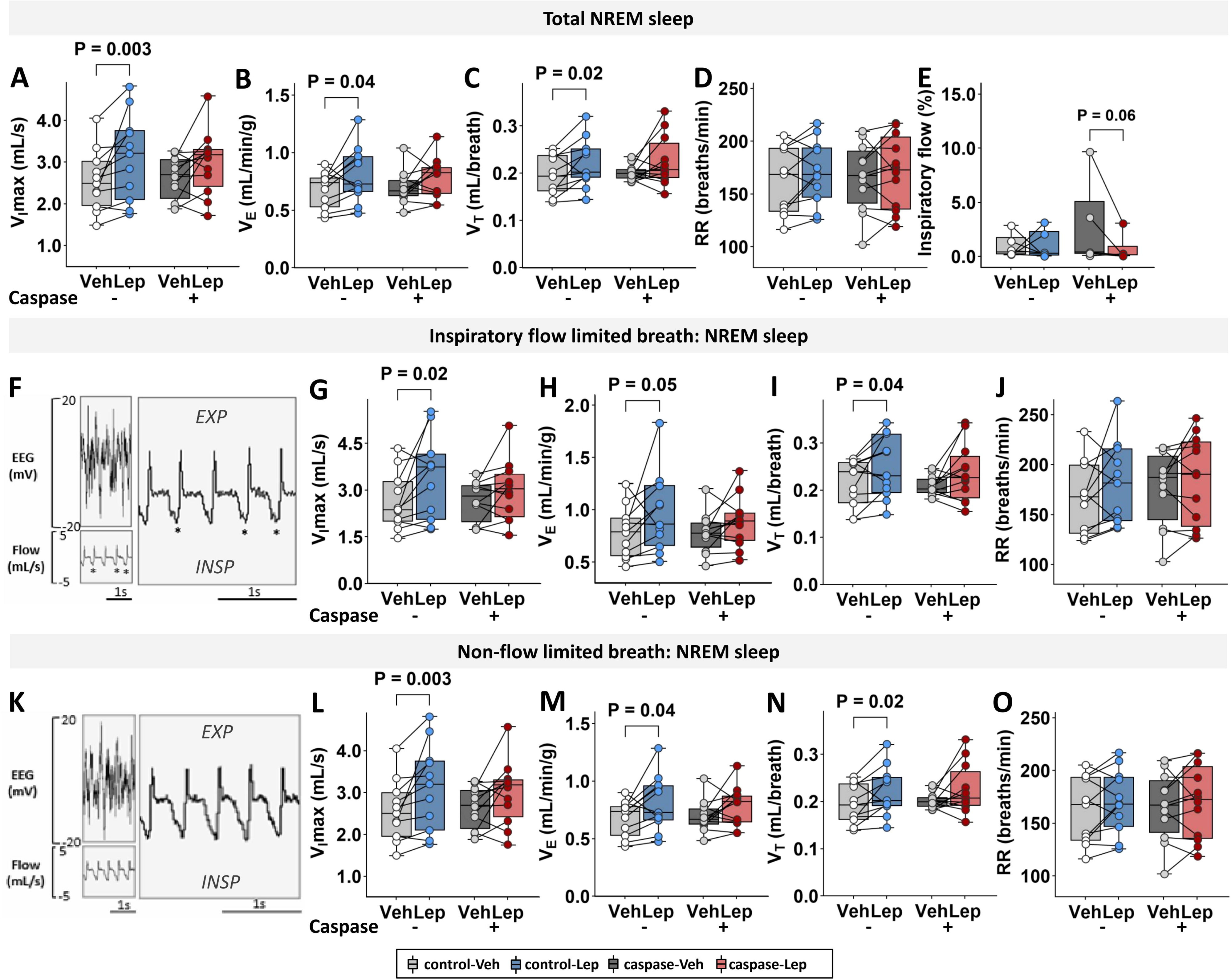
Breathing during NREM sleep. The deletion of the 5-HT neurons projecting to the MR prevented the respiratory effects of intranasal leptin during NREM sleep. The effect of leptin and 5-HT neuron deletion on **(A)** maximal inspiratory flow (V_I max_), **(B)** minute ventilation (V_E_), **(C)** tidal volume (V_T_), **(D)** respiratory rate (RR) and **(E)** % of inspiratory flow limited breaths. **(F)** Representative recording of the inspiratory flow limited breaths (*). Inspiration is down; expiration is up. The analysis of inspiratory flow limited breathing: **(G)** V_I max_, **(H)** V_E_, **(I)** V_T_, and **(J)** RR. **(K)** Representative recording of the non-flow limited breathing (*). **(L)** V_I max_, **(M)** V_E_, **(N)** V_T_, and **(O)** RR. Male control virus, n=11; male caspase virus, n=12. Data are plotted using boxplots (median ± 1.5*interquartile range). Statistical analyses were performed using the Wilcoxon matched-pairs signed rank test or Mann-Whitney test. Exact P values are shown in the figures.

### The effect of intranasal leptin on breathing during REM sleep and the role of the 5-HT neurons projecting to the XII MN

In control mice, leptin increased V_E_, but not V_I max_ during both flow limited and non-flow limited breathing, although the effect was mediated by an increase in respiratory rate during flow limited breathing and by an increase in V_T_ during non-flow limited breathing **(Fig. 6**). Similar to NREM sleep, ablation of 5-HT neurons projecting to the XII MN abolished effects of leptin on V_E_ and brought up the leptin-induced decrease in the frequency of flow limited breaths (**Fig. 6E**). Thus, leptin stimulates breathing during REM sleep *via* serotonergic innervation of the XII MN.

**Figure 6.**
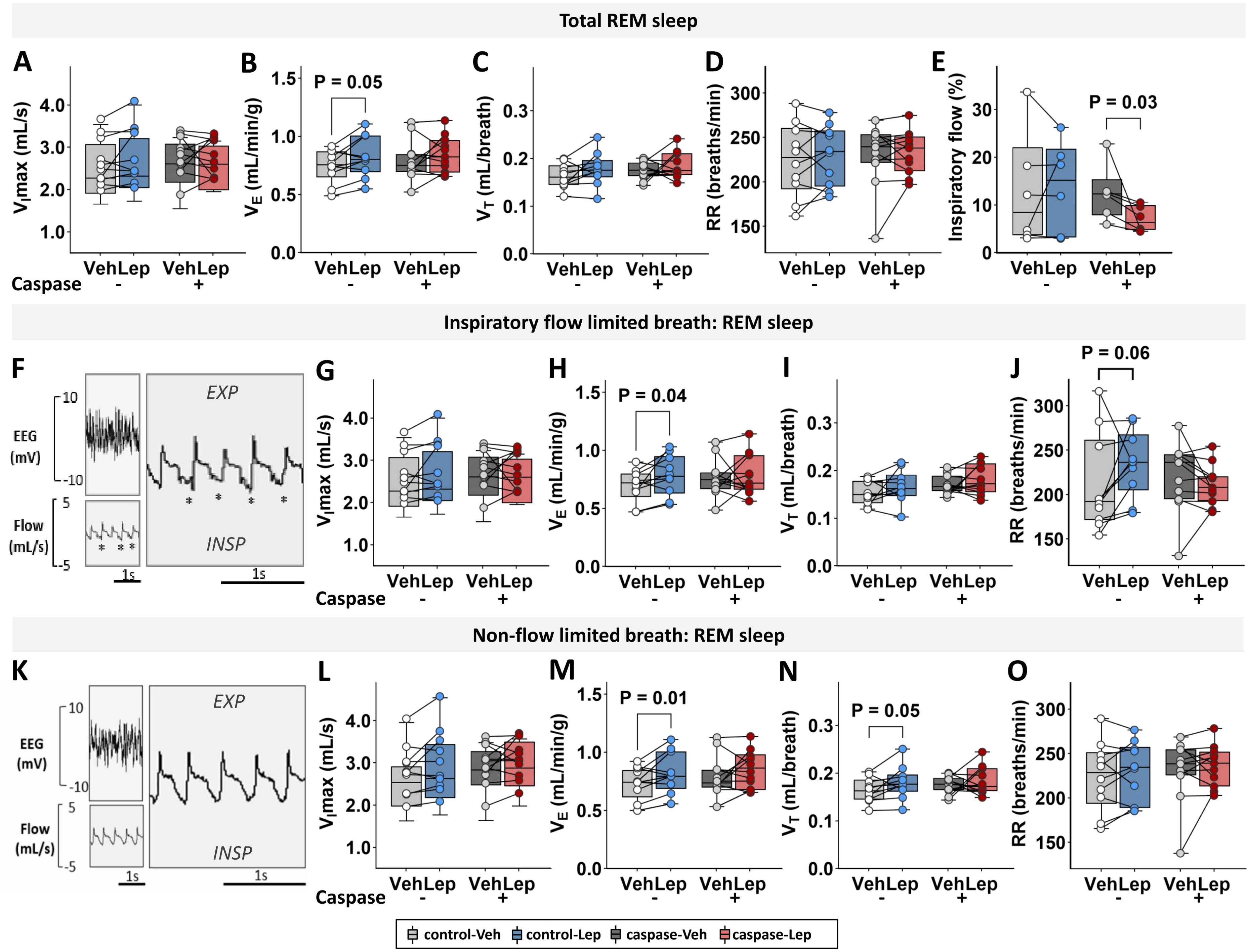
Breathing during REM sleep. The deletion of the 5-HT neurons projecting to the MR prevented the respiratory effects of intranasal leptin during REM sleep. The effect of leptin and 5-HT neuron deletion on **(A)** maximal inspiratory flow (V_I max_), **(B)** minute ventilation (V_E_), **(C)** tidal volume (V_T_), **(D)** respiratory rate (RR) and **(E)** % of inspiratory flow limited breaths. **(F)** Representative recording of the inspiratory flow limited breaths (*). Inspiration is down; expiration is up. The analysis of inspiratory flow limited breathing: **(G)** V_I max_, **(H)** V_E_, **(I)** V_T_, and **(J)** RR. **(K)** Representative recording of the non-flow limited breathing (*). **(L)** V_I max_, **(M)** V_E_, **(N)** V_T_, and **(O)** RR. Male control virus, n=11; male caspase virus, n=12. Data are plotted using boxplots (median ± 1.5*interquartile range). Statistical analyses were performed using the Wilcoxon matched-pairs signed rank test or Mann-Whitney test. Exact P values are shown in the figures.

### The effect of intranasal leptin on GG muscle activity and the role of the 5-HT neurons projecting to the XII MN

As expected, GG muscle activity increased by CO_2_ and progressively decreased in normocapnia from wakefulness to NREM and REM sleep in control mice, providing quality assurance^25^ (**Figure S5**). Intranasal leptin significantly augmented GG activity under normocapnic and hypercapnic conditions during wakefulness (**Figure 7A, D, E**) and under normocapnic conditions during NREM sleep (**Figure 7B, E**), whereas no significant effect was observed during REM sleep (**Figure 7C**). Ablation of 5-HT neurons projecting to the XII MN with *FlpO*-dependent caspase abolished the effects of leptin. Thus, leptin increases GG activity under normocapnic and hypercapnic conditions during wake and NREM sleep *via* serotonergic innervation of the XII MN.

**Figure 7.**
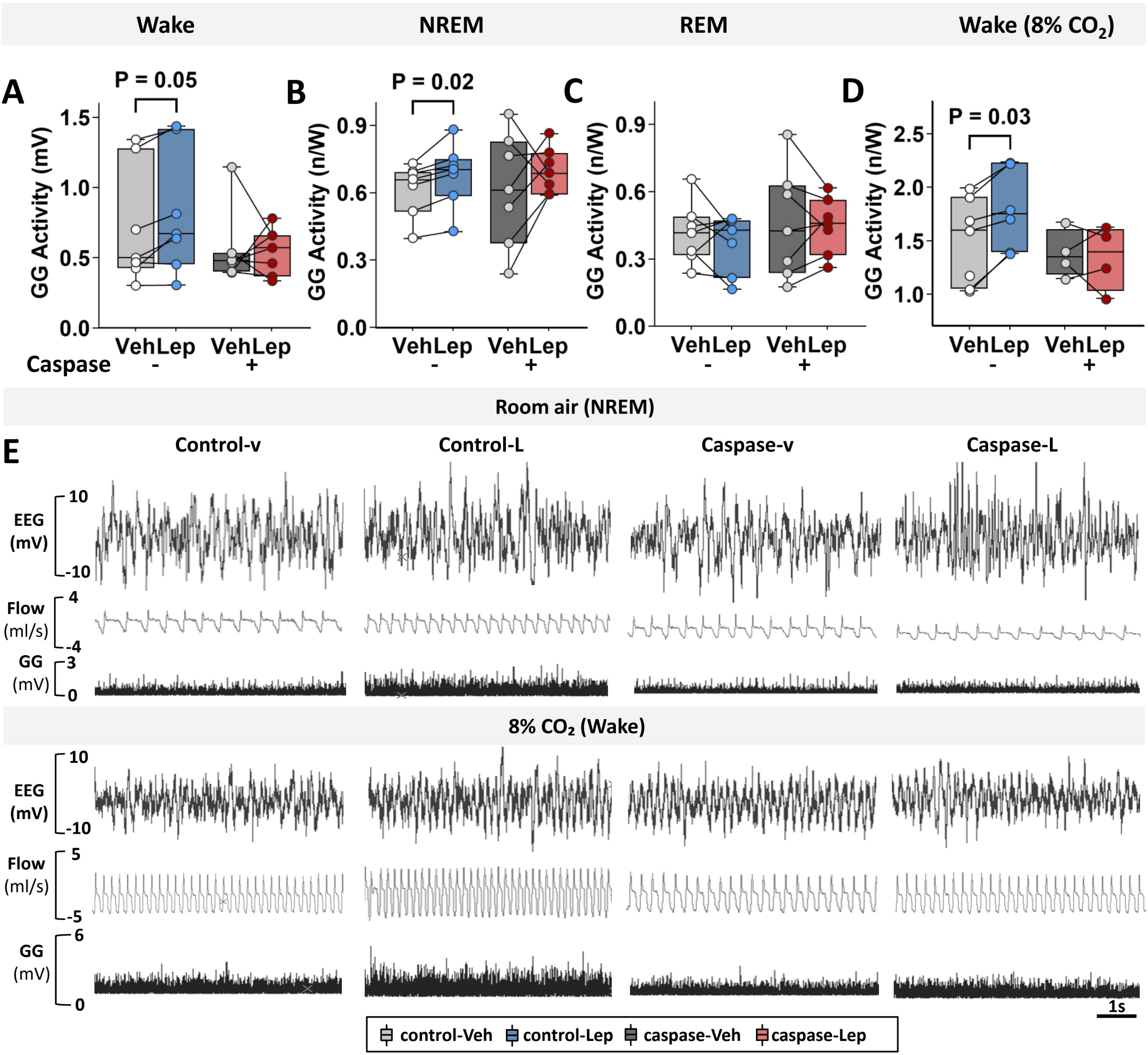
Genioglossus (GG) muscle activity across sleep/wake stages. Intranasal leptin augmented GG muscle activity during NREM sleep in the room air and enhanced GG activity during 8% CO_2_ exposure. Leptin induced enhancement of GG activity was abolished by elimination of 5-HT neurons with retrograde AAV caspase. **GG** activity in **(A)** Wake, **(B)** NREM, **(C)** REM sleep, and **(D)** 8% CO_2_ exposure. **(E)** The representative figure of GG activity changes on each group. Male control virus, n=4-7; male caspase virus, n=4-7. Data are plotted using boxplots (median ± 1.5*interquartile range). Statistical analyses were performed using the Wilcoxon matched-pairs signed rank test or Mann-Whitney test. Exact P values are shown in the figures. n/W, values normalized to quiet wakefulness in room air. Lep, leptin; Veh, vehicle; control-Veh, control-vehicle; control-Lep, control-leptin; caspase-Veh, caspase-vehicle; caspase-Lep, caspase-leptin.

### LEPR^b^-expressing DMH neurons project to the medullary raphe (MR)

We have previously shown that LEPR^b^ DMH neurons project to the serotonergic DR^49^. Here, we injected AAV-Floxed-YFP virus into the dorsomedial hypothalamus (DMH) of *LEPR^b^-Cr*e mice and examined projections to the MR (**Figure 8A**). There were numerous projections of the LEPR^b^ DMH neurons (green) to the 5-HT neurons (red) of the raphe obscurus (RO) and raphe pallidus (RPA) (**Figure 8B**). Thus, the LEPR^b^ DMH neurons have numerous direct projections to the MR.

**Figure 8.**
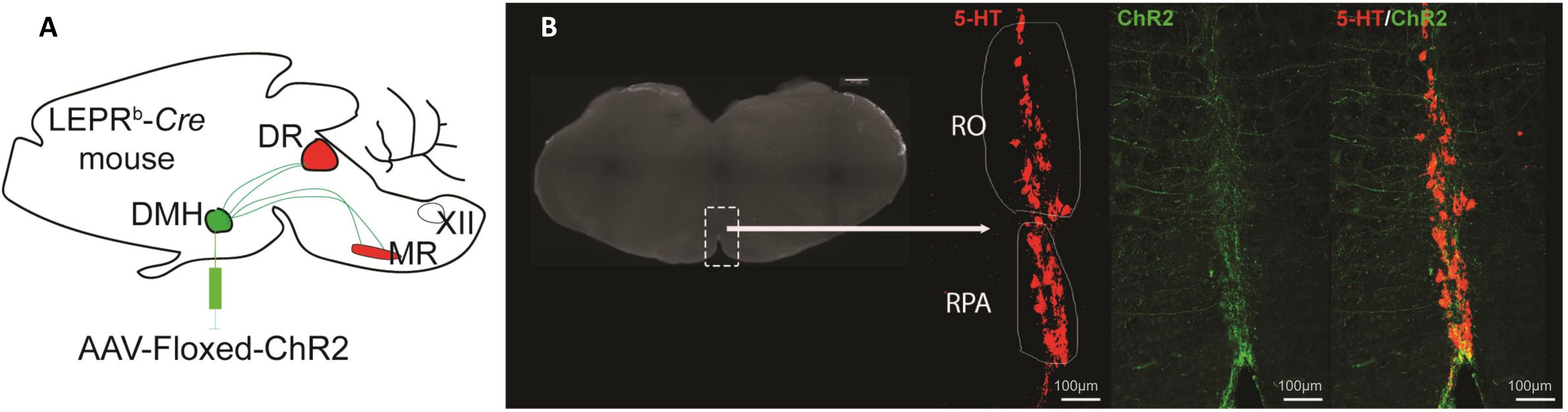
LEPR^b^+DMH neurons show projections to the medullary raphe (MR) nuclei. Top: Left: the schematic of the YFP-tagged viral injection to DMH (green) with projections to the MR (red); Center: the slice of the medulla with the box denoting MR. Panels showing 5-HT and LEPR^b^-ChR2-YFP fibers colocalize resulting in the orange color (right). RO, raphe obscurus; RPA, raphe pallidus. Scale bars 100 µm.

## Discussion

We have previously shown that IN leptin abolished upper airway obstruction during sleep and augmented hypercapnic chemoreflex in DIO mice treating SDB^43,49^. We have also shown that leptin acts on LEPR^b^+ DMH neurons^48,49^, which project to the DR serotonergic nucleus in the brainstem^49^. However, it remained unknown if the respiratory effects of leptin maybe are mediated *via* 5-HT innervation of the hypoglossal nucleus. To our knowledge, we provide the first evidence that IN leptin increases genioglossus muscle tone during NREM sleep and in response to hypercapnia, which could be an important mechanism of the therapeutic effect of IN leptin in OHS^45^. The main finding of our paper is that elimination of the 5-HT neurons innervating the hypoglossal motoneurons abolished leptin-induced augmentation of the HCVR, the leptin-induced increase in upper airway muscle tone during sleep, and the leptin-induced increases in minute ventilation during NREM and REM sleep.

Transfection of *Sert-flp* mice with retrograde AAV, which cannot cross the synaptic cleft and harbors *FlpO*-dependent tracer or caspase, showed that 5-HT neurons of medullary raphe (MR) and, specifically, raphe obscurus (RO) and raphe pallidus (RPA) monosynaptically innervate hypoglossal motoneurons **(Fig. 1; Fig. S1).** Two subpopulations of 5-HT MR neurons (RO) modulating respiratory chemoreflex have been previously identified by Dymecki’s lab: the *Egr2-Pet1* neurons mediate chemoreception, but do not innervate respiratory motonuclei ^71^, whereas *Tac1-Pet1* neurons respond to CO_2_ and project directly to respiratory motor neurons including the hypoglossal nucleus^72^. In contrast to our MR findings, we did not find any evidence of direct connections between the DR and XII nuclei, although the presence of GFP+ fibers in DR suggests that the MR neurons projecting to the hypoglossal motoneurons also project to DR. The abundance of 5-HT neurons in the MR projecting to XII motor neurons, together with their complete elimination using retrograde AAV carrying a 5-HT-specific caspase, enabled us to define the role of serotonergic innervation of the upper airway in the respiratory effects of leptin.

Our study provided definitive evidence that leptin stimulates the genioglossus muscle and attenuates upper airway obstruction during sleep *via* serotonergic pathways. Our previous human and animal work showed that upper airway obstruction is defined by inspiratory airflow limitation, which is characterized by an early inspiratory plateau in airflow at a maximum level (V_I max_) while effort continues to increase^44,73–75^. We have linked the effects of leptin to activation of the XII MN^43,47^, which dilate the pharynx by stimulating the main tongue protrudor, the GG muscle^12,22,76^. Here we have shown that IN leptin increases both GG activity and V_I max_ during flow limited obstructed breaths in awake and sleeping mice. Furthermore, GG muscle activity predicted V_I max_ in REM sleep (**Fig. 9B)**, when inspiratory flow limitation is the most prevalent (**Fig. 6E**), but not in NREM sleep (**Fig. 9A**) when flow limited breaths are seldom **(Fig. 5E**). The effects of leptin on upper airway muscle activity and patency were abolished by elimination of 5-HT MR neurons suggesting that 5-HT MR (predominantly RO and RPA) neurons mediate effects of leptin on upper airway obstruction during sleep.

**Figure 9.**
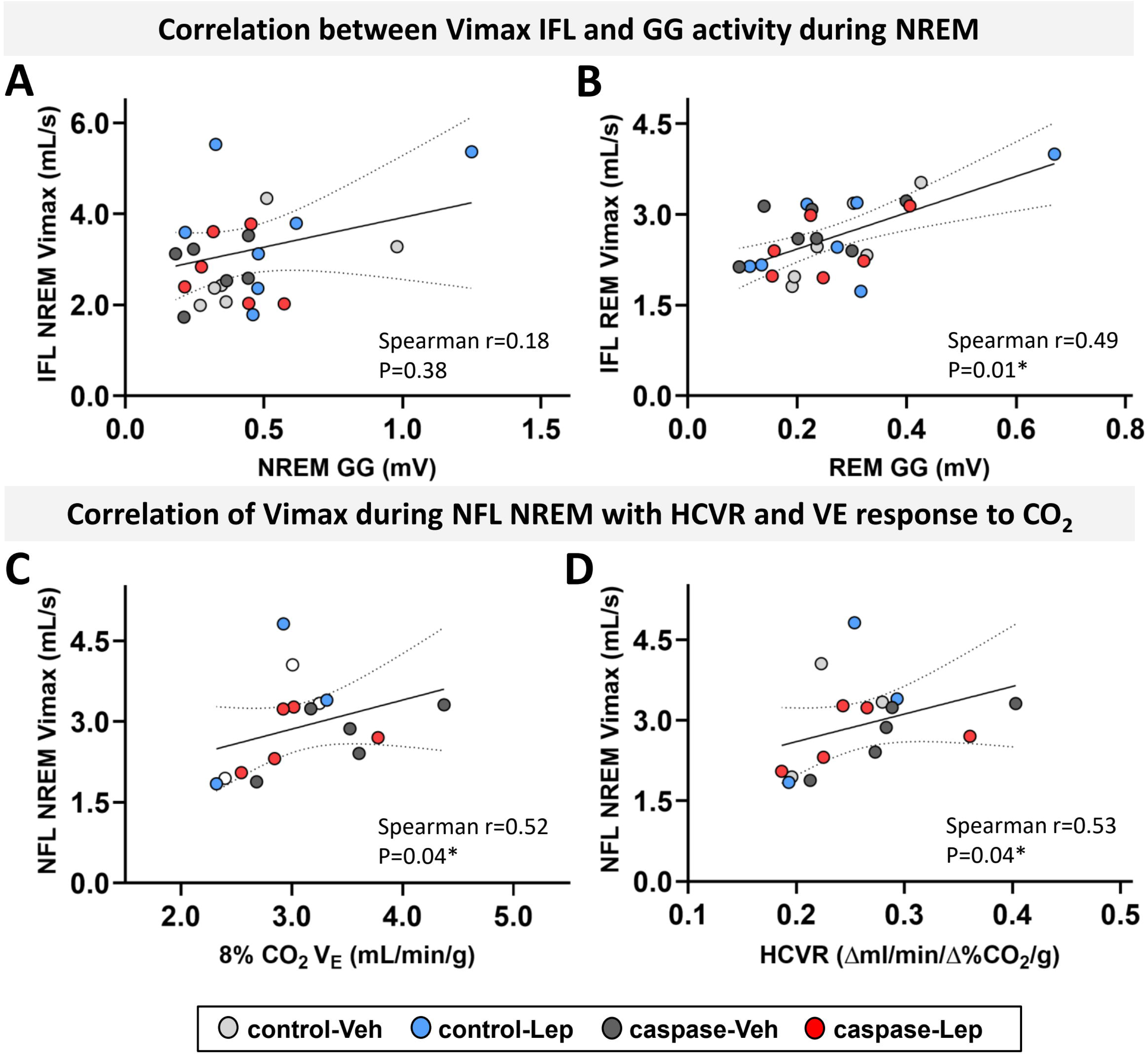
Physiological correlates of maximal inspiratory flow (V_I max_) during flow limited and non-flow limited breathing. Correlations between genioglossus (GG) muscle activity and V_I max_ during flow-limited breathing during NREM **(A)** and REM **(B)** sleep. Correlations between **(C)** hypercapnic minute ventilation (V_E_) in awake mice and **(D)** the hypercapnic ventilatory response (HCVR) and V_I max_ during non-flow limited breathing during NREM sleep. Scatter plot with fitted linear trend line. Association assessed using two tailed Spearman’s correlation analysis. Exact P values and “r” are shown in the figures.

**Figure 10.**
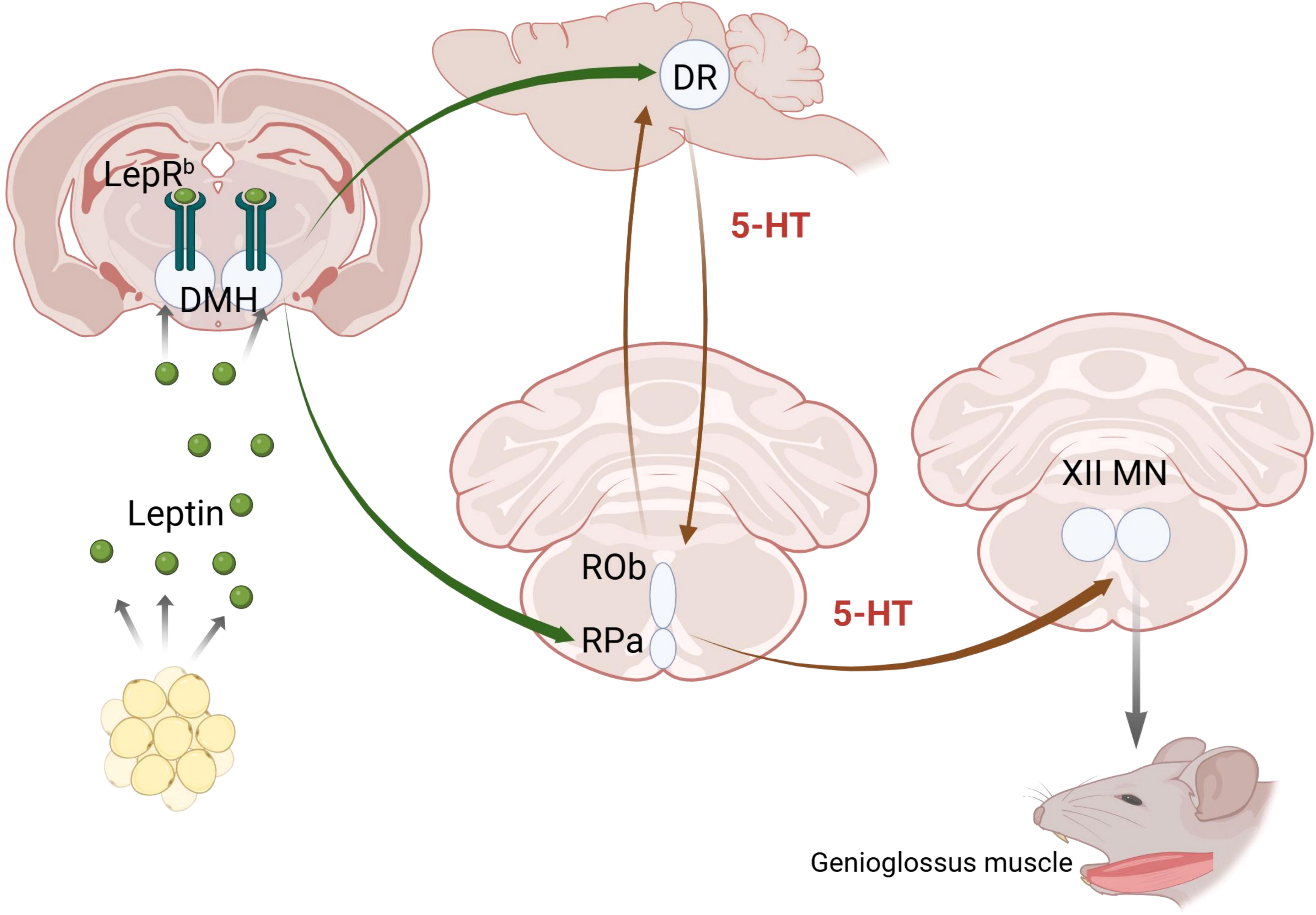
Leptin regulates breathing during sleep by modulating serotonergic innervation of the upper airways. DMH, dorsomedial hypothalamus; DR, dorsal raphe nucleus; LEPR^b^, leptin receptor; RO, raphe obscurus; RPA, raphe pallidus; XII MN, hypoglossal motor nucleus; the hypercapnic ventilatory response (HCVR).

We reproduced previous work by our group and others showing that leptin enhances the HCVR in male *ob/ob* and DIO mice^34,49,77^ and that these effects are mediated via the 5-HT axis^49^. Here we provide novel evidence that 5-HT neurons innervating the XII nucleus are also responsible for the leptin’s effect on hypercapnic sensitivity. One possibility is that the same neurons, possibly *Tac1-Pet1* neurons^72^, directly mediate both upper airway and ventilatory control effects. It is also conceivable that leptin-induced relief of upper airway obstruction resets PaCO_2_ during sleep and improves CO_2_ sensitivity, similarly to patients with OHS in whom upper airway obstruction is eliminated by CPAP treatment^78–80^. In the absence of airway obstruction, i.e. during non-flow limited breaths, ventilation is controlled by metabolic rate and hypercapnic sensitivity. Neither leptin nor 5-HT neurons affected VCO_2_ (**Figure S4**). HCVR in awake mice predicted V_I max_ during non-flow limited breathing in NREM and REM sleep **(Fig. 9C&D**) and the effects of leptin on V_I max_ and V_E_ during sleep were abolished by caspase **(Figs. 5&6).** All of the above suggests that 5-HT neurons projecting to the XII nucleus mediate effects of leptin not only on hypercapnic sensitivity, but also on ventilation during sleep.

We have previously shown that LEPR^b^ is scantily expressed in the DR and MR^49^. We have also shown that respiratory effects of leptin are mediated *via* LEPR^b^+ DMH neurons projecting to the DR^49^. Our present data suggests that leptin treats SDB by acting on 5-HT MR neurons. LEPR^b^+ DMH neurons may stimulate hypoglossal motoneurons polysynaptically *via* the DR -> MR pathway (**Figure 8**). Alternatively, LEPR^b^+ DMH neurons directly project to 5-HT RO and RPA (**Fig. 8**) and may activate XII neurons and GG muscle without the DR involvement. Prior literature suggests that DR is not directly implicated in hypercapnic sensitivity, but is involved in hypercapnic arousals^81–83^. Leptin did not disrupt sleep nor decrease arousal latency **(Figs. 3&4).** In fact, leptin increased hypercapnic arousal latency **(Figure 3B)** and decreased the number of spontaneous arousals from sleep in control mice (**Figure 4**), probably due to its beneficial effects on the upper airway, since these effects were abolished by elimination of 5-HT neurons projecting to the hypoglossal nucleus (**Figures 3B&4**). LEPR^b^+ DMH neurons may simultaneously project to DR and MR, and caspase-mediated elimination of DR-projecting LEPR^b^+ DMH neurons could abolish effects of leptin because these neurons also project to the 5-HT MR neurons. Taken together, our previous^49^ and current data imply that leptin acts on LEPR^b^+ DMH neurons treating OHS *via* serotonergic stimulation of the upper airway muscles mediated by 5-HT MR neurons (**Figure 8**).

Although leptin greatly decreased severity of airway obstruction measured by V_I max_, it did not affect the frequency of flow limited breaths in control mice. In contrast, leptin significantly decreased the frequency of flow limited breaths upon elimination of the 5-HT MR neurons projecting to the XII nucleus, which suggests that leptin may also improve airway patency *via* non-serotonergic pathways **(Figure 4E&5E)**. Stimulation of LEPR^b^+ NTS neurons did not affect genioglossus muscle activity or airway patency in DIO mice^84^. However, chemogenetic stimulation of LEPR^b^ + NTS neurons projecting to the DMH and lateral parabrachial nucleus attenuated central apneas in mice^85^, which we did not observe in our study **(Figure S3)**. Thus, non-5-HT mechanisms and pathways may also be implicated in leptin’s action on control of breathing and upper airway function.

Finally, our study showed significant sex differences. Neither leptin nor caspase-mediated elimination of 5-HT innervation of the hypoglossal nucleus had any effect on the HCVR or arousal latency in female mice. We have previously shown that female mice are protected against respiratory effects of DIO and do not develop hypoventilation, HCVR suppression or significant SDB. DIO females exhibit higher ventilation at baseline and at hypercapnic conditions, compared to DIO males^46,49,86^, and do not respond to leptin^49,70^.

Our study had several limitations. First, we did not perform sleep studies in female DIO mice, because HCVR data and our previous publications^49,86^ suggest that female DIO are protected against OHS and respiratory leptin resistance and neither leptin nor 5-HT will be effective, probably due to a ceiling effect. Second, our study was limited to studying the role of 5-HT neurons monosynaptically projecting to the XII nucleus in the effects of leptin. Other populations of 5-HT neurons as well as noradrenergic, cholinergic, glutamatergic and other neurons may also be involved in the leptin pathways. Third, leptin did not affect the apnea-hypopnea index in DIO mice. However, apneas in mice are nearly exclusively central. Our data indicate that the main effect of leptin on SDB is elimination of upper airway obstruction. Fourth, studies showed the efficacy of glucagon-like peptide 1 receptor (GLP1R) agonists^87^, a combination of noradrenergic and anticholinergic agents^29,88^ as well as carbon anhydrase inhibitors^89,90^ for human OSA, whereas serotonergic agents were ineffective^91^. Our pre-clinical study suggests that serotonergic agents may be effective in leptin resistant subjects with OHS.

## Conclusion

Our pre-clinical study in an established murine model of OHS showed that leptin treats OHS by stimulating upper airway muscles *via* 5-HT MR neurons projecting to the hypoglossal nucleus.

## AUTHOR CONTRIBUTIONS

**Conception and Design:** VYP, MRA and DM; **Data Acquisition:** DD, MAR, JLD, MKS, XW; **Data Analysis & Interpretation:** DD, CRW, VYP; **Drafting and Critical Revision:** DD, DM and VYP; **Final Approval:** All authors have reviewed and approved the final version.

## CONFLICTS OF INTEREST

The authors declare no conflict of interests

## FUNDING

The study has been funded by NIH R01HL128970, R01 HL174409, and NIH S10OD032420

## ACKNOWLEDGEMENTS

We are grateful to Drs. Anastas Popratiloff and Cheryl Clarkson-Paredes for their expert assistance with confocal imaging. BioRender.com has been used to create Figures 1 and 10.

## Notes

### Competing Interest Statement

The authors have declared no competing interest.

