## supplemental figures for "Leptin alleviates obesity hypoventilation *via* serotonergic pathways"

**SUPPLEMENTARY FIGURES**

**Figure S1.** Representative images illustrating direct projections of hypoglossal neurons (yellow fluorescent protein, green color) to the serotonergic neurons (tryptophan hydroxylase, red color) of the raphe obscurus (RO) and raphe pallidus (RPA). Merged orange color denotes colocalization.

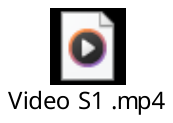

**Figure S2. Delta power of NREM sleep after intranasal leptin or vehicle treatment has been normalized to delta power of NREM sleep prior to the intervention during the same study** (Relative delta power)**.** Deletion of MR 5-HT neurons following injection of control or caspase virus into XII N did not affect **(A)** total delta power of NREM sleep. **(B)** delta power measured across sequential hourly time intervals from 10:00 AM to 5:00 PM under four experimental groups. Data points represent mean values for each time interval.

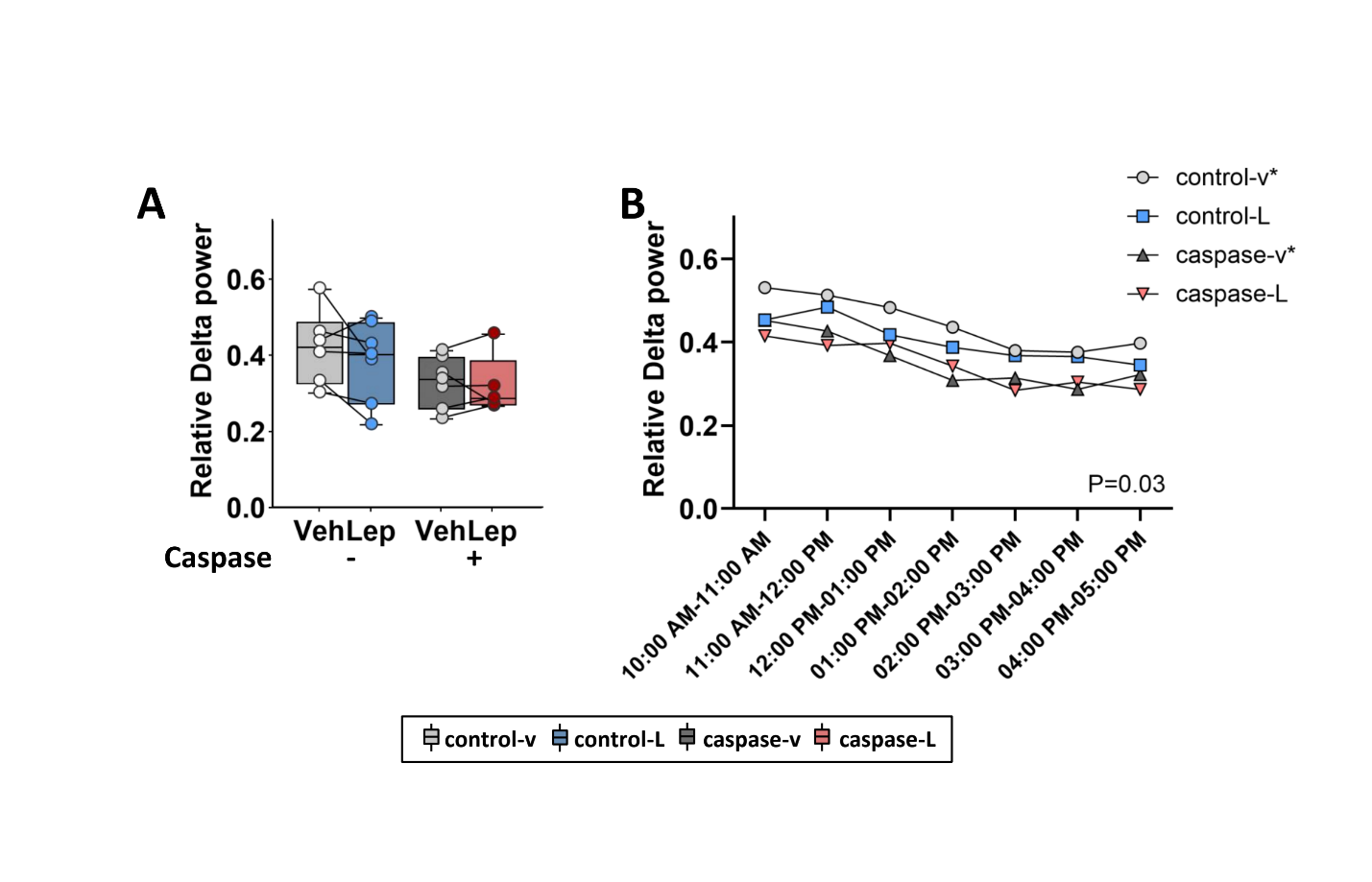

**Figure S3. Intranasal leptin and deletion of MR 5-HT neurons following injection of control and caspase virus into XII N had no effect** on **(A)** the apnea index or **(B)** the sigh index.

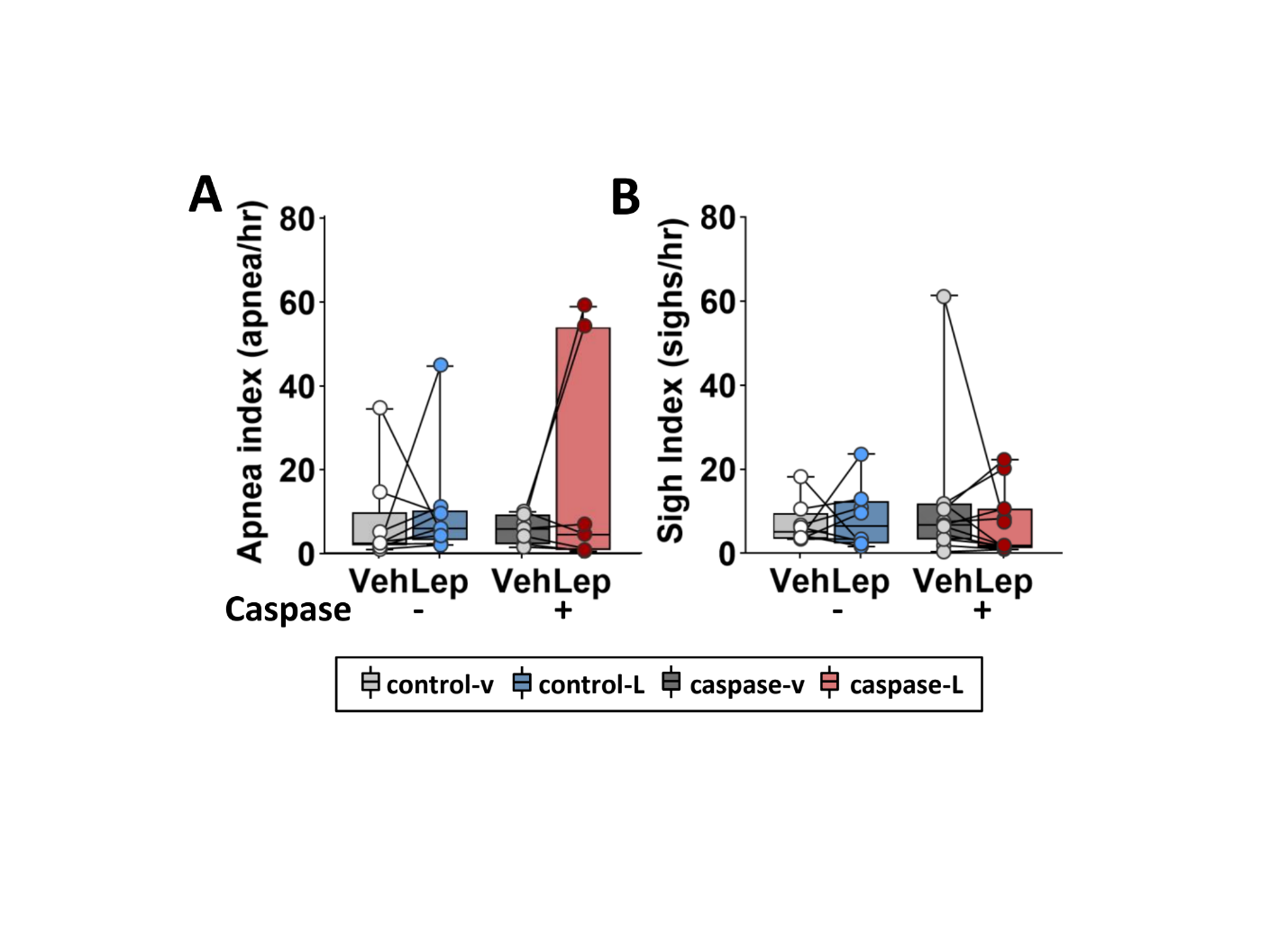

**Figure S4.** Metabolic measurements over 24 hours after intranasal leptin vs vehicle treatment in mice transfected with control or 5-HT dependent caspase harboring viruses. **(A-D)** the light phase and **(E-H)** the dark. **(A and E)** Total oxygen consumption (VO_2_), **(B and F)** total carbon dioxide production (VCO_2_), **(C and G)** respiratory exchange ratio (RER), **(D and H)** motor activity. Minute ventilation (V_E_) adjusted for metabolic indexes during **(I and J)** light and **(K and L)** dark phase in the control and caspase groups after intranasal leptin. Control virus, n=6; Caspase virus, n=7. Data are plotted using boxplots (median ± 1.5*interquartile range). Statistical analyses were performed using the Wilcoxon matched-pairs signed rank test or Mann-Whitney test. Exact P values are shown in the figures.
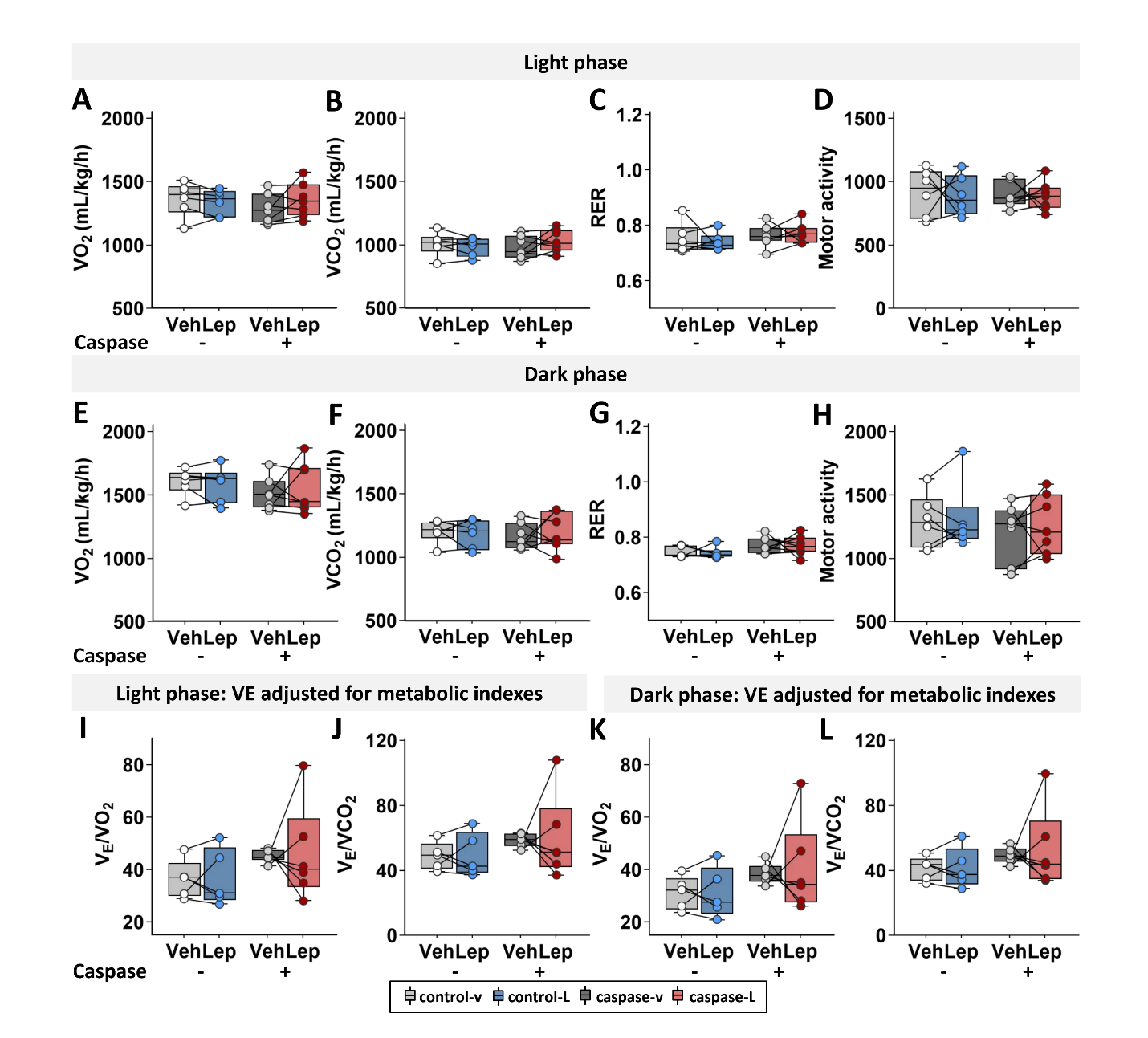

**Figure S5**. **The quality control of the genioglossus (GG) lead implantation.** Changes genioglossus (GG) muscle activity across sleep states. **(A)** GG activity progressively decreased from wakefulness to NREM and REM sleep, with significant reductions between wakefulness and NREM, and between NREM and REM sleep. **(B)** GG activity during exposure to room air and CO_2_. GG activity increased during CO_2_ exposure compared with room air. Individual data points are connected by lines, and colored circles represent individual subjects. Values are expressed in arbitrary units (a.u.). *p ≤ 0.05 and **p < 0.01 using the Wilcoxon matched-pairs signed rank test or Mann-Whitney test.

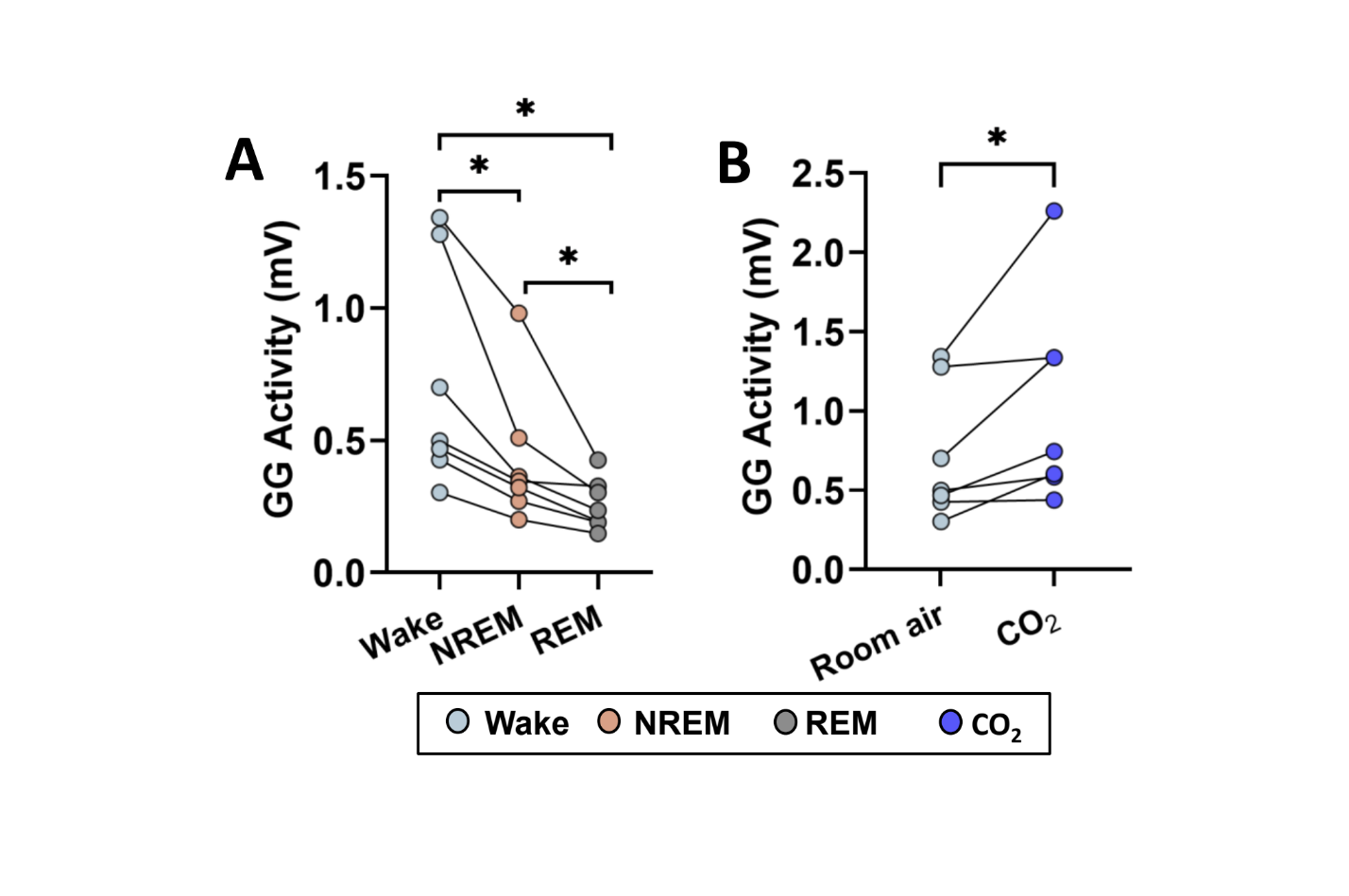
